# General features of cognition gene polymorphism patterns in marine fish

**DOI:** 10.1101/2025.11.22.689885

**Authors:** Zhizhou Zhang, Shuaiyu Zhang, Yongdong Xu, Changlu Guo

## Abstract

A growing body of evidence suggests that the cognitive abilities of fish are far more advanced than traditionally believed. They are not mere “automata” driven by instinct, but complex organisms capable of learning, remembering, solving simple problems, engaging in social interactions, and even deception. This situation has prompted us to explore genes related to cognitive functions in the fish genome, examining the pattern status of a set of SNVs (single nucleotide variations) in their cognitive genes relative to the human genome sequence. This study investigates the locus status of 1,824 SNVs from 33 human cognitive genes across the whole genomes of 90 fish species. Based on this, preliminary findings indicate that fish exhibit approximately four patterns of cognitive gene polymorphism. One of these patterns includes at least two coelacanth/lungfish samples, which directly point to the evolutionary direction of the ancient human sd1 sample (an ancient human hair sample from Sudan, Africa).

---

Our previous research suggested that the polymorphism patterns of cognitive genes in ancient humans appear to be directly derived from marine fish rather than from other primates [1-2]. This implies that the framework for human linguistic and cognitive genes might have begun to be constructed as early as during the marine fish stage. This point is also easy to understand, as even the most complex biological functions (such as language and cognition) generally evolve step by step from simple to complex [3-6]. It is akin to how a complex high-rise building must be constructed gradually from what appears to be a simple foundation and design framework. In prior studies, we used approximately 500 whole-genome samples, covering major animal groups, including over 100 ancient human samples, among which the oldest ancient human sample is sd1 (an ancient human hair sample from Sudan, Africa; **Table S1**). Therefore, one of the questions in this study is which marine fish samples are evolutionarily closest to sd1; however, the main question of this study is how many groups the polymorphism patterns of cognitive genes in marine fish can roughly be categorized into, and which of these groups shows an evolutionary trajectory toward the ancient human sd1.

To address the aforementioned questions, it is essential to first prepare sufficiently extensive and high-quality data. The primary data consists of SNV loci sourced from 33 cognitive genes (**Table S2**). Several previous studies utilized only 462 SNVs (**Table S3**), a number that is decidedly insufficient. Due to computational limitations in our laboratory, even this study is confined to using only 1,824 SNVs (**Table S4**). Corresponding research based on 12,000 or more SNVs will only become feasible several months from now, as our current software, HAXI07plus03 (version 7.3) [7-9], for searching tens of thousands or more SNV sequences within whole-genome sequence files, is not yet fast enough. Completing the search across several hundred FASTQ whole-genome files (general file size: 20-200 GB) indeed requires several months or even longer. Moreover, to obtain preliminary results promptly, this study utilizes representative samples from various animal groups, totaling 261 samples (**Table S5**). These include 90 fish samples, 26 reptile samples, 6 amphibian samples, 8 rodent samples, 11 bird samples, 27 Laurasiatheria samples, 33 Primate samples, 35 ancient human samples, and 25 other samples. Therefore, this study is based on 261 whole-genome samples and 1,824 SNVs from 33 cognitive genes.The sequence status of these modern human SNVs across various species’ genomes was determined by searching the 261 whole-genome samples using the HAXI07plus03 software. (All SNV loci and their flanking sequences were extracted from the respective gene sequences using R packages. The distribution of each SNV across the 33 cognitive gene sequences is illustrated in **Figure S1**). The actual search employed 27-base-long strings (the SNV locus plus 13 bases on each side) for exact matching. Results with no exact match were recorded as “0”. Consequently, the search result for each locus has 16 possible states: A, T, G, C, AT, AG, AC, TG, TC, GC, ATG, ATC, AGC, TGC, ATGC, and 0.The resulting SNV data based on the 261 samples are presented in **Table S5**, and the taxonomic data for these samples are provided in **Table S6**.

First, we performed PCoA (Principal Coordinate Analysis) on the 1,824 SNV dataset from the 261 whole-genome samples, as shown in Figure 1. PCoA arranges samples in a low-dimensional space based on a distance matrix (such as Euclidean distance, Bray-Curtis distance, etc.), so that the distances in the low-dimensional space approximate the distances in the original space as closely as possible. In the analysis of SNV data, PCoA can visualize the similarities between samples using any distance matrix (e.g., Hamming distance, Jaccard distance, etc.). It may effectively reveal evolutionary relationships or population structure among the samples. Figure 1 shows that sd1 is the ancient human sample closest to marine fish and amphibians. Several other ancient human samples relatively close to marine fish and amphibians include nd2, nd7, jm1, dg2, etc., from Eurasia. If the ancient human population represented by sd1 is actually the ancestor of samples like nd2, nd7, jm1, and dg2, this would be consistent with the current conclusion of human origins in Africa. However, the relationship between the origin of modern *Homo sapiens* and the aforementioned samples remains unclear.

**Figure 1.**
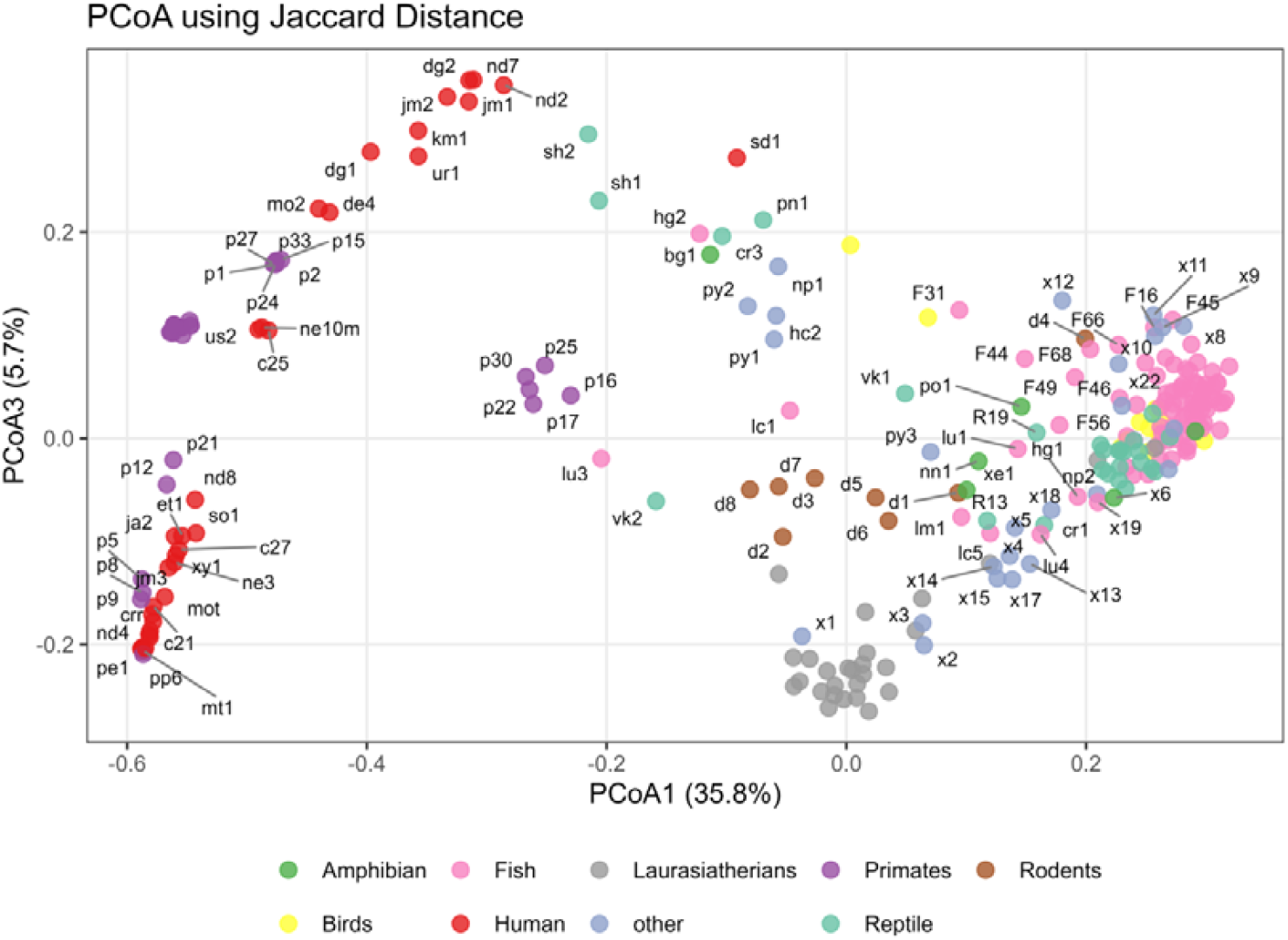
PCoA results of 1,824 SNVs from all 261 samples.

To more clearly observe the distribution characteristics of cognitive gene polymorphism patterns in fish, we retained only 151 samples (**Table S7**: 5 amphibians, 11 birds, 88 fish, 1 ancient human, 21 others, 23 reptiles, 1 rodent; taxonomic data provided in **Table S8**) based on Figure 1 and performed further PCoA analysis. The result is shown in Figure 2. In Figure 2, the fish samples are basically distributed across four regions. The majority of samples are concentrated in the lower right corner, six samples are in the upper right corner, seven samples are located in the middle lower left area, and two samples are positioned in the middle left. These four regions essentially represent four distinct patterns of cognitive gene polymorphism.

**Figure 2.**
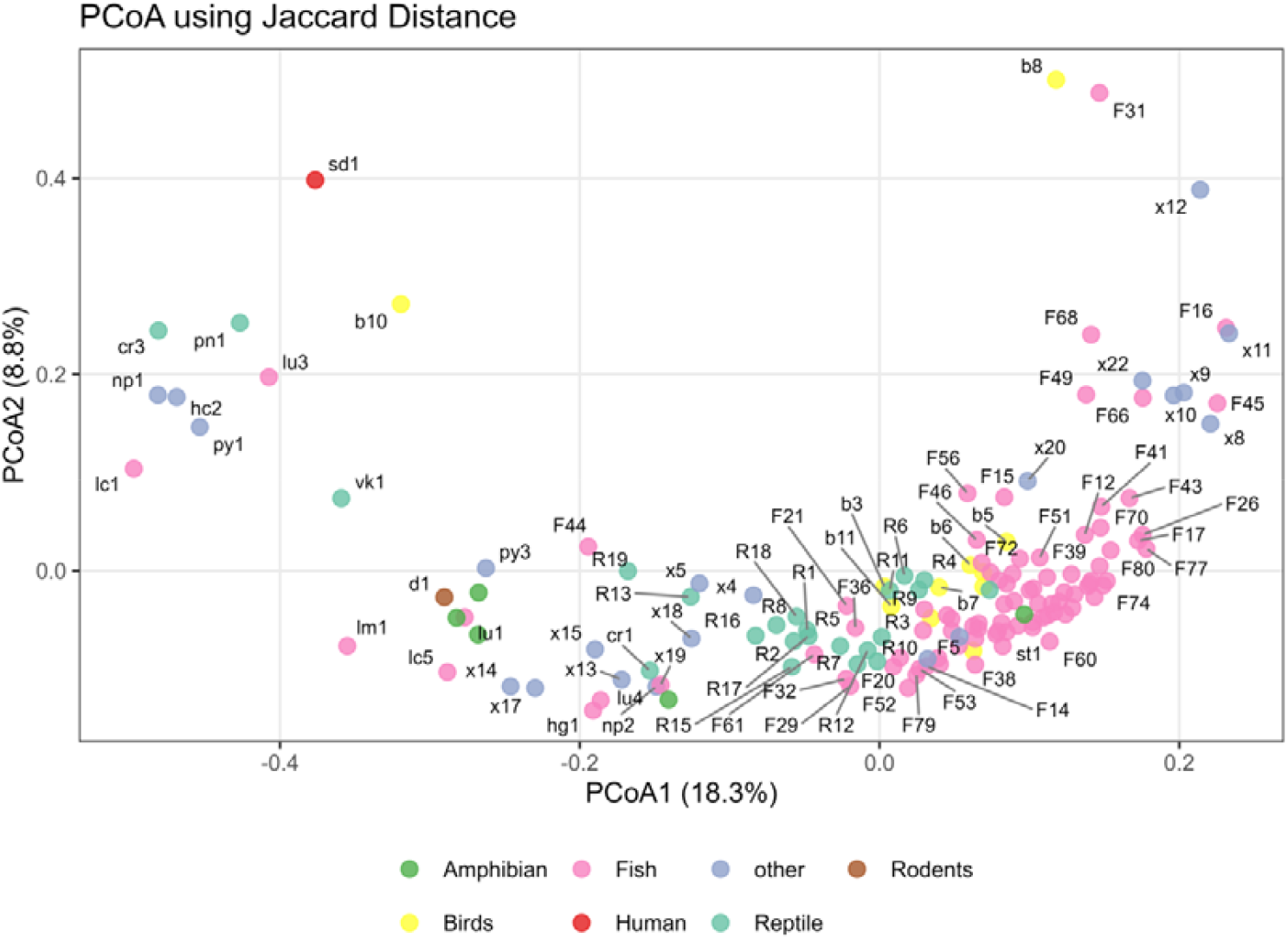
PCoA results of 1,824 SNVs from the 151 samples. This figure allows for a closer observation of the fish, amphibian, or reptile samples closest to the ancient human sample sd1, and further identifies several patterns of cognitive gene polymorphism in fish.

Several computational methods for measuring complex string similarity (such as Nei’s genetic distance, Jaccard distance, Levenshtein distance, etc.) can also provide information on the patterns of cognitive gene polymorphism in fish [10]. The upper part of Figure 3 shows the Nei’s genetic distance curve, while the lower part displays the corresponding first-order derivative curve. Since there are two distinct inflection points in Panel A of Figure 3, and the polymorphism patterns on either side of these inflection points differ significantly, the polymorphism patterns in fish can be roughly categorized into four types based on this observation, which aligns with the results shown in Figure 2. The first inflection point is near the lungfish samples lu3 and lu4, and the second inflection point is near F72 and F74, with the lungfish sample lu1 located very close to this second inflection point. The position where the rate of change in the polymorphism pattern is fastest in Panel A of Figure 3 is near the sample py1 (platypus) (as indicated by the first peak in the derivative curve), and the coelacanth sample lc1 is relatively close to py1. Figure 4 overlays the results of seven similarity calculations (Cosine distance, Euclidean distance, Nei’s genetic distance, TANIMOTO distance, DICE distance, Jaccard distance, and Rogers distance [11-12]), demonstrating that the cognitive gene polymorphism patterns in fish samples (although intermixed with samples from other groups) indeed underwent two significant changes (with two distinct inflection points).

**Figure 3.**
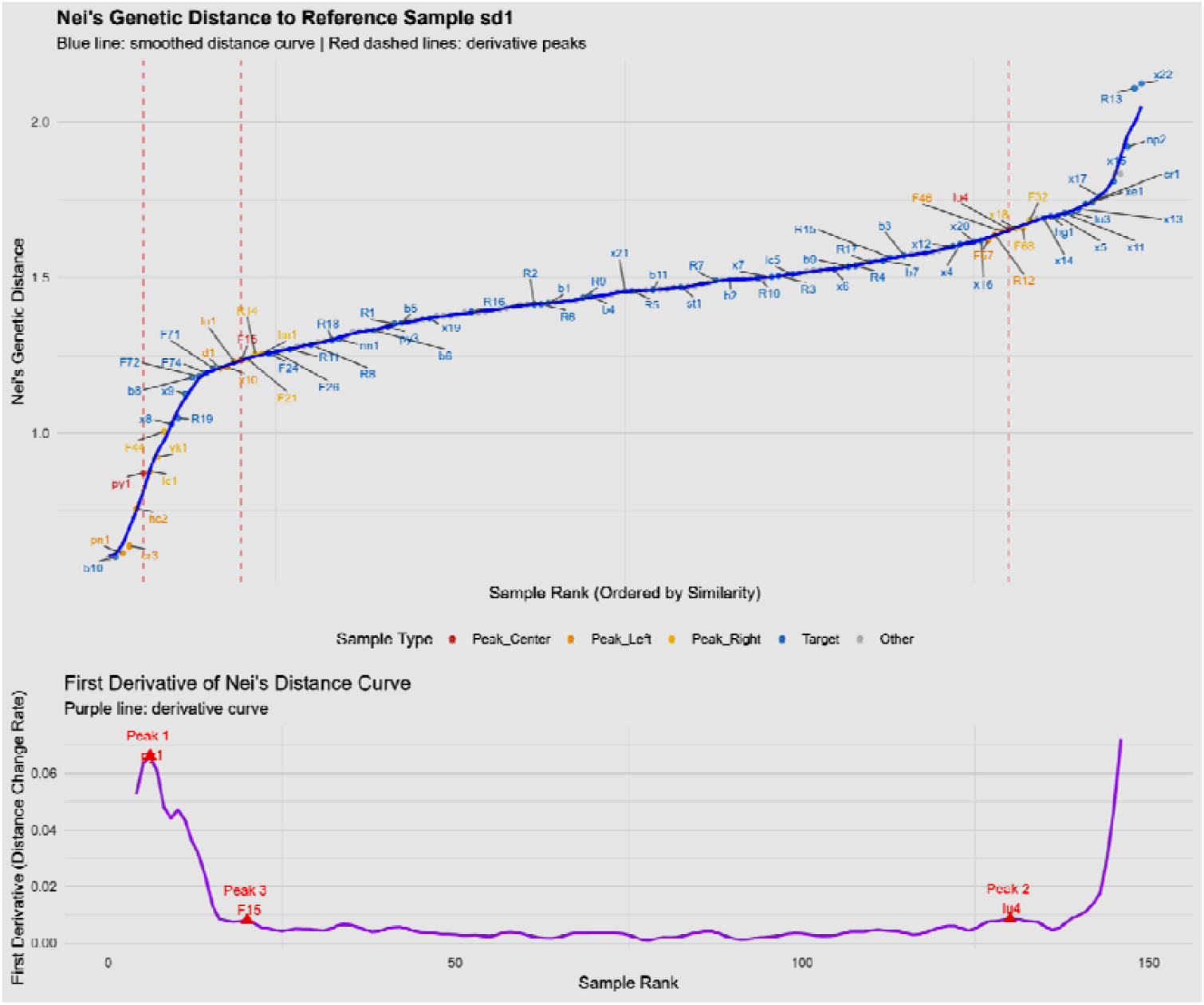
The Nei’s genetic distance plot reveals inflection points where significant changes in cognitive gene polymorphism patterns occurred in fish (relative to the ancient human sample sd1).

**Figure 4.**
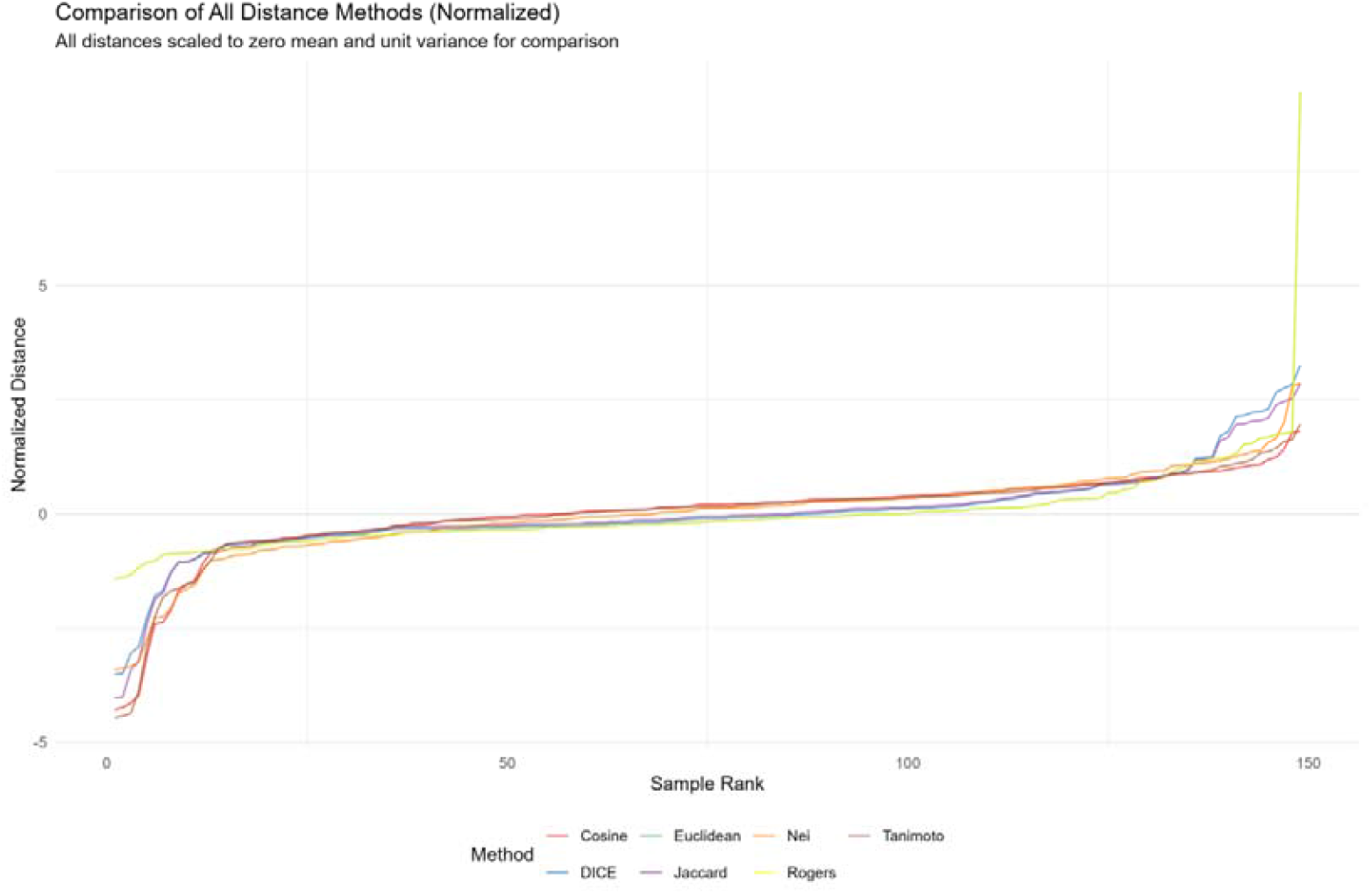
Overlay plot of seven similarity curves (relative to the ancient human sample sd1). Similarity values are arranged from small to large along the horizontal axis.

Figure 5 displays the sample RANK positions (arranged in ascending order of similarity) corresponding to the four largest peaks on the first-derivative curves of ten similarity measures. Among these, eight curves show three derivative peaks within the first 20 samples, while the remaining two curves exhibit four derivative peaks—two located at the extreme ends of the sample RANK and two within the first 30 samples. Given the generally lower reliability of peaks at the extremes of the RANK, the samples corresponding to the third derivative peak in the eight curves demonstrate relatively higher credibility. These likely represent critical junctures in the evolutionary trajectory of cognitive gene polymorphism patterns, warranting further detailed investigation in future studies. Two samples, lm1 (Indonesian coelacanth *Latimeria menadoensis*) and vk1 (Komodo dragon), appear most frequently near the third derivative peak across the eight curves. The first-derivative peak data for all ten similarity measures are provided in **Supplementary Document-1**.

**Figure 5.**
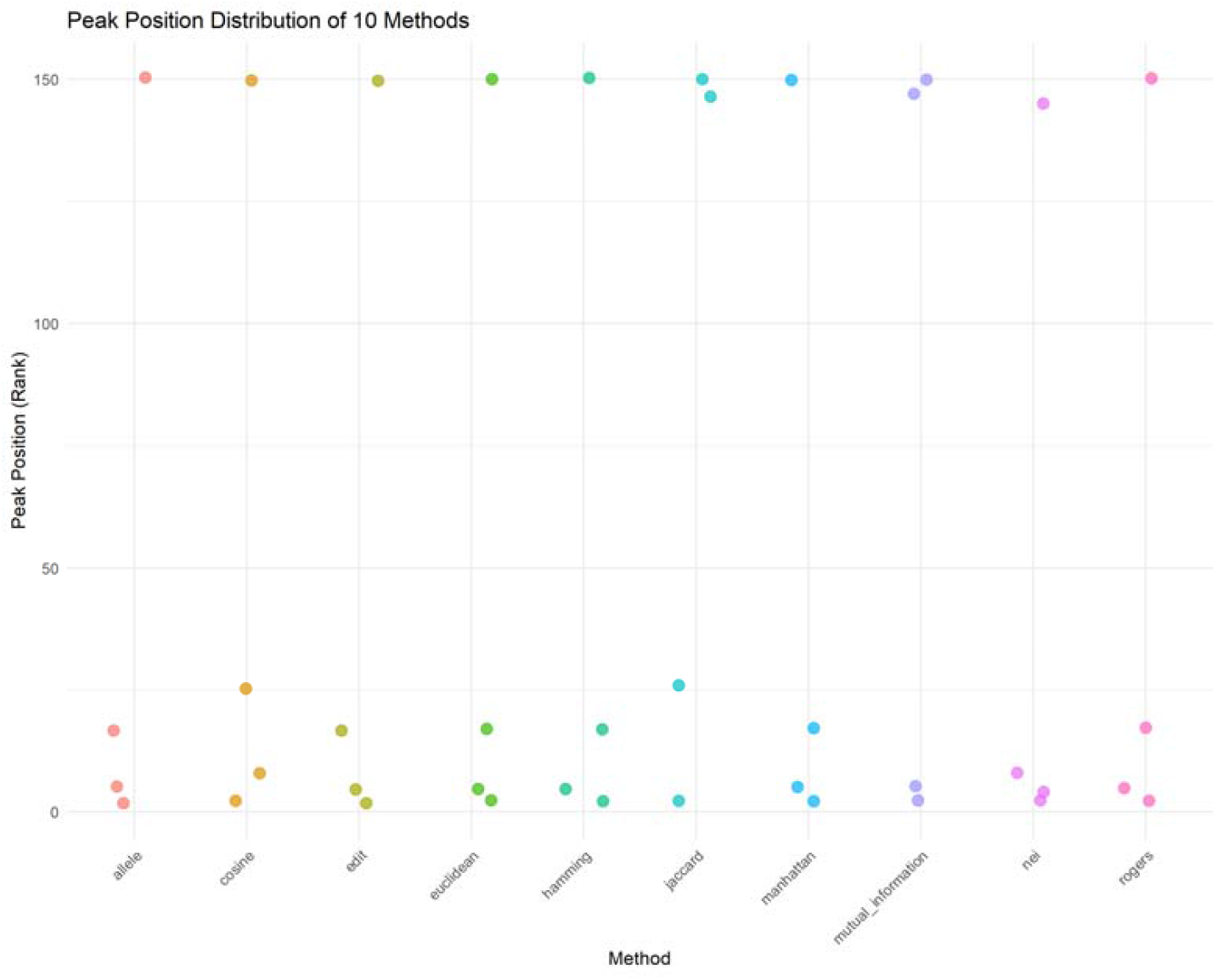
Distribution of the four peak values from the derivative curves of ten similarity measures within the sample RANK . The R code for all similarity curve-related calculations in this study can be obtained from the corresponding author upon request.

The phylogenetic tree in Figure 6 was constructed using samples from ancient humans, fish, amphibians, reptiles, and birds. Although this tree does not readily reveal the approximately four patterns of cognitive gene polymorphism in fish, it clearly shows that the fish samples closest to ancient humans are actually lc1 (*Latimeria chalumnae* Tanzania), lu1 (South American lungfish), hg1 (Japan Yaizu brown hagfish *Eptatretus atami*), and F44 (Mexican tetra) [13-18]. Solely from the perspective of cognitive gene polymorphism evolution, F44 may represent the common ancestor of sd1, lc1, lu1, and hg1.

**Figure 6.**
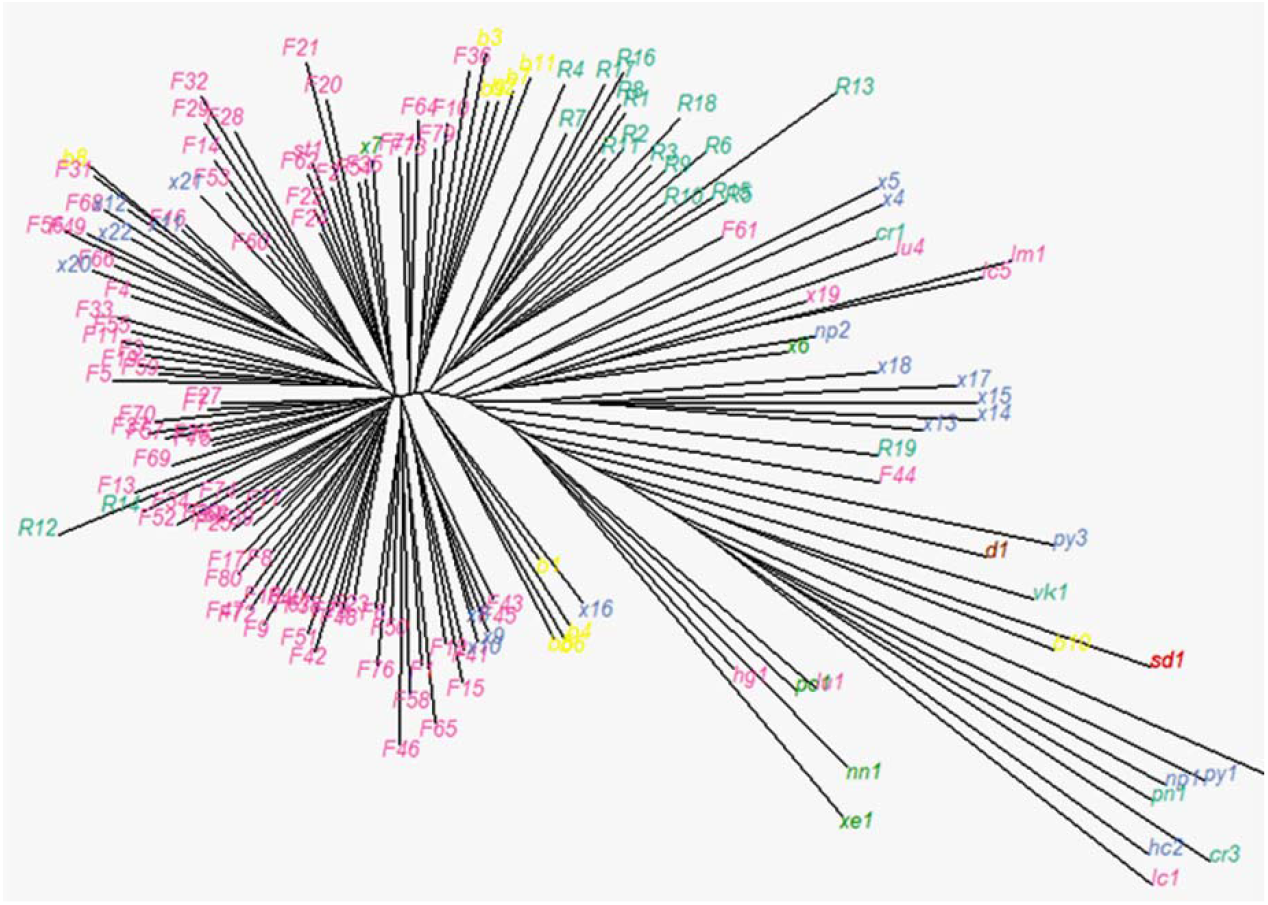
The phylogenetic tree reveals more important details regarding the evolution of cognitive gene polymorphism patterns in fish. Pink represents fish samples, red represents ancient human samples, green represents amphibians, and yellow represents birds. The remaining samples belong to reptiles.

## Summary

This study employed 261 whole-genome sequence samples (90 fish, 26 reptiles, 6 amphibians, 8 rodents, 11 birds, 27 Laurasiatheria, 33 primates, 35 ancient humans, and 25 other samples) to examine the locus status of 1,824 SNVs from 33 cognition-related genes across these specimens. Through cluster analysis and a series of DNA string similarity calculations, we discovered that the fish samples exhibit approximately four types of cognitive gene polymorphism patterns. One of these patterns, which includes specimens such as the coelacanth lc1 (*Latimeria chalumnae* Tanzania) and lu1 (South American lungfish), points toward the evolutionary direction that gave rise to the ancient human sd1 (the ancient human hair sample from Sudan, Africa).

## Acknowledgments

This study was supported by a State Language Commission Research Grant (YB135-117), Association of Chinese Graduate Education Grant (B-2017Y0505-079), and National Research Center for Foreign Language Education Grant (ZGWYJYJJ10A042).

