## supplementary tables and figures for "General features of cognition gene polymorphism patterns in marine fish": Table S2 33 selected human cognition genes.docx

Table S2 Selected human cognition genes in this study^[a-g]^

|  | Gene | Cognition gene | Language-related | Function or Compromised ability (example) when mutated |
| --- | --- | --- | --- | --- |
| 1 | ARHGAP11B | √ |  | Hominin-specific development and evolutionary expansion of the brain neocortex |
| 2 | ASPM | √ | √ | Associated with [microcephaly](https://www.malacards.org/card/microcephaly_5_primary_autosomal_recessive) |
| 3 | MCPH1 | √ | √ | Associated with [microcephaly , primary, autosomal recessive](https://www.malacards.org/card/microcephaly_1_primary_autosomal_recessive) and [lymphatic malformation](https://www.malacards.org/card/lymphatic_malformation_10); |
| 4 | CHRM2 | √ |  | A nervous system gene associated with depression disorder |
| 5 | IGF2R | √ |  | Insulin-like growth factor gene associated with behavior/neurological phenotype |
| 6 | Snap25 | √ |  | A gene associated with neurotransmitter release |
| 7 | Fads2 | √ |  | A member of the fatty acid desaturase，associated with craniofacial abnormalities |
| 8 | Dab1 | √ |  | A gene linked with nervous system development |
| 9 | NBPF8 | √ |  | A gene associated with macrocephaly, autism, schizophrenia, cognitive disability |
| 10 | HAR1A | √ |  | A gene whose expression levels associated with memory and cognitive abilities |
| 11 | GNB5 | √ |  | Associated with language delay and cognitive Impairment |
| 12 | NRXN1 | √ |  | Neurexin 1required for efficient neurotransmission and formation of synaptic contacts |
| 13 | DCC | √ |  | Associated with impaired intellectual development |
| 14 | GRID2 | √ |  | Predominant excitatory neurotransmitter receptors in the mammalian brain |
| 15 | EP300 | √ |  | Associated with rare neurological diseases and impairment of intellectual development |
| 16 | KMT2D | √ |  | Lysine Methyltransferase 2D, associated with intellectual disability and eye diseases |
| 17 | NOTCH2NL | √ |  | Neural progenitor proliferation and evolutionary expansion of the brain neocortex |
| 18 | THSD7B | √ |  | Associated with eye diseases/neuronal diseases |
| 19 | FOXP1 | √ | √ | Expressive language |
| 20 | FOXP2 | √ | √ | Speech |
| 21 | CNTNAP2 | √ | √ | Early language development |
| 22 | TPK1 | √ | √ | Syntactic and lexical ability |
| 23 | DCDC2 | √ | √ | Reading, dyslexia |
| 24 | KIAA0319 | √ | √ | Reading, dyslexia |
| 25 | TM4SF20 | √ | √ | Language delay; communication disorder |
| 26 | FLNC | √ | √ | Reading, language |
| 27 | ATP2C2 | √ | √ | Memory |
| 28 | ROBO1 | √ | √ | Phonological buffer |
| 29 | ROBO2 | √ | √ | Expressive vocabulary |
| 30 | CMIP | √ | √ | Reading, memory |
| 31 | NFXL1 | √ | √ | Speech |
| 32 | SRGAP2 | √ | √ | Vocal learning, vital for cortical neuron development |
| 33 | SRGAP2C | √ | √ | Modifier of cortical connectivity in the human brain |

[a] Li M, Zhang W, Zhou X. 2020. Identification of genes involved in the evolution of human intelligence through combination of inter-species and intra-species genetic variations. PeerJ 8:e8912

[b] Natalia A. Goriounova and Huibert D. Mansvelder. Genes, Cells and Brain Areas ofIntelligence.Frontiers in Human Neuroscience,2019

[c] Savage et al. Genome-wide association meta-analysis in 269,867 individuals identifies new genetic and functional links to intelligence.Nat Genet. 2018 July; 50(7):912-919.

[d] Suzanne Sniekers et al. Genome-wide association meta-analysis of 78,308 individuals identifies new loci and genes influencing human intelligence.Nat Genet.2017 July; 49(7):1107-1112.

[e] Wei Xia, Zhizhou Zhang. Language gene polymorphism pattern survey provided important information for education context in human evolution. biorxiv 2022

[f] Lei Shi et al. Regional selection of the brain size regulating gene CASC5 provides new insight into human brain evolution. Hum Genet. 2017 Feb;136(2):193-204.

[g] Tattersall I. Endocranial volumes and human evolutionF1000Research 2023, 12:565 <https://doi.org/10.12688/f1000research.131636.1>
