## Supplementary figures and images for "General features of cognition gene polymorphism patterns in marine fish"

### fig s1 1824 SNVs distribution.png

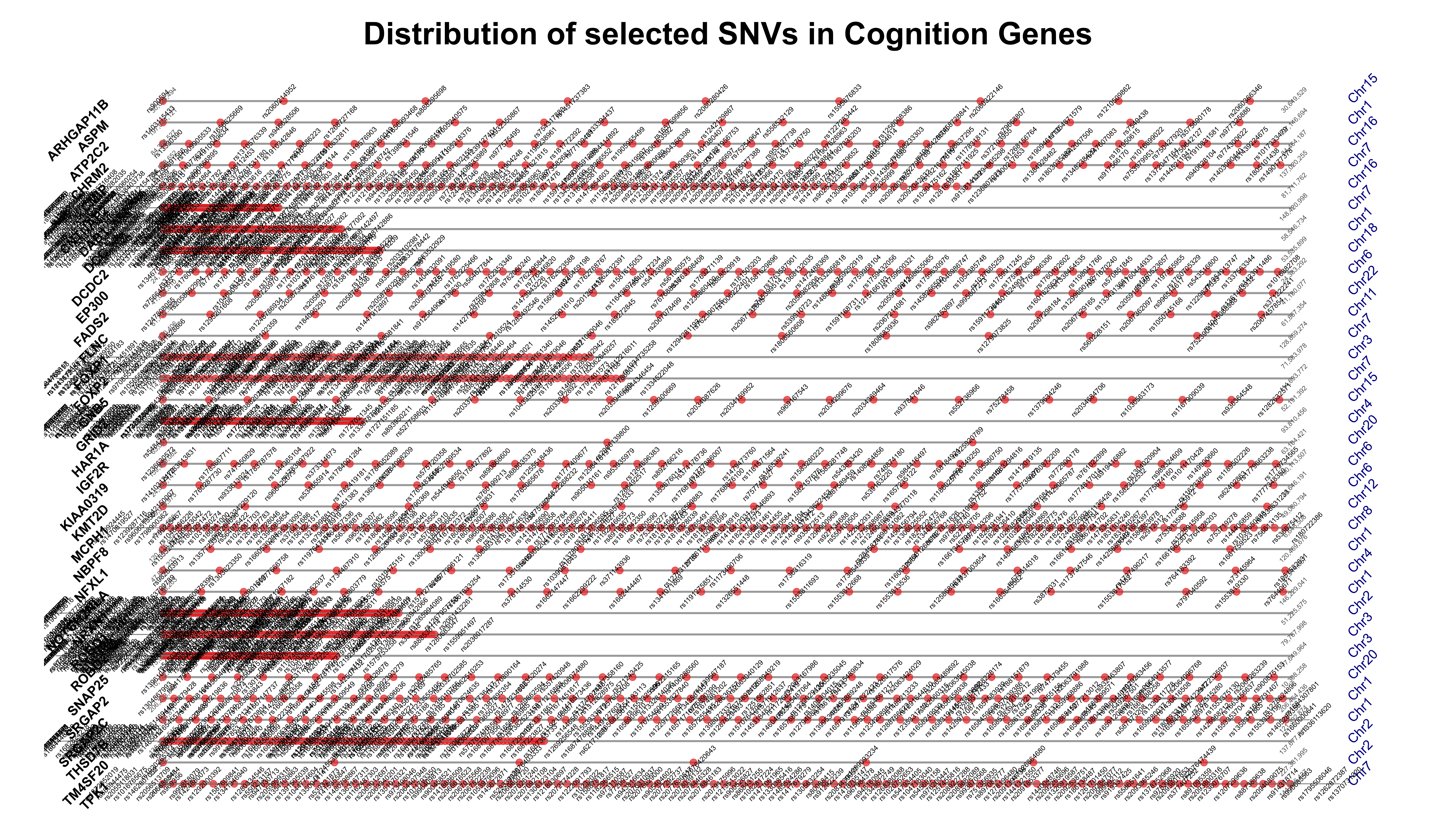
